# RAGE protein: COPD biomarker and case study for variant pathogenicity prediction

**DOI:** 10.1101/2025.08.04.668362

**Authors:** JE Dawson, RP Bowler

## Abstract

The structural biology of a protein is key to understanding its mechanism and plays a significant role in predicting the effects of clinical variants. New advances in pathogenicity predictors have improved their quality but interpreting the results in light of a protein’s structure and functional can still be challenging. We use the Receptor for Advanced Glycosylation End-products (RAGE), which is involved in triggering inflammatory pathways during Chronic Obstructive Pulmonary Disease (COPD), as a case study to demonstrate how to utilize publicly available structure and missense variant databases, such as ClinVar and the Genomic Aggregation database (gnomAD), as well as the pathogenicity predictor AlphaMissense. Common missense RAGE variant G82S (rs2070600) has an observed allele frequency of 0.056 in gnomAD and is predicted to be benign, but is known to have clinical consequences in COPD, AD, and CV, which illustrates the need to careful interpretation of results. Additionally, missense variants for a set of COPD-related proteins were utilized to evaluate the performance of pathogenicity predictors SIFT, FATHMM, Poly-Pred2, MutPred2, and AlphaMissense.

## Introduction

The structural biology of a protein is key to understanding its mechanism and plays a significant role in predicting the effects of clinical variants. For example, the receptor advanced glycosylation end-product (RAGE) protein’s domain structure and oligomeric state are involved in activating the receptor. RAGE is encoded by the advanced glycosylation end-products (*AGER*) gene, is a leading protein risk biomarker for chronic obstructive pulmonary disease (COPD) and emphysema susceptibility and progression^1^. COPD and emphysema is caused by irritant exposure (smoke, pollution) leading to shortness of breath (airflow). RAGE is a transmembrane, multi-ligand receptor, which is expressed on many cells^2^, most predominantly in the lung epithelium^3^. Smoke, inflammation, and aging cause the release of DAMP (damage-associated molecular pattern) ligands (**Figure 1**). Ligand-binding and RAGE oligomerization^4-8^ triggers inflammation *via* the NF-κB pathway and leads to IL-6 and other cytokine secretion^9-11^. Both RAGE and its ligands can be up-regulated by the inflammatory response. RAGE ligand signaling can be blocked with soluble RAGE, which is formed either through shedding of the ectodomain of RAGE (sRAGE) or through expression of an alternate splice variant (esRAGE).

**Figure 1.**
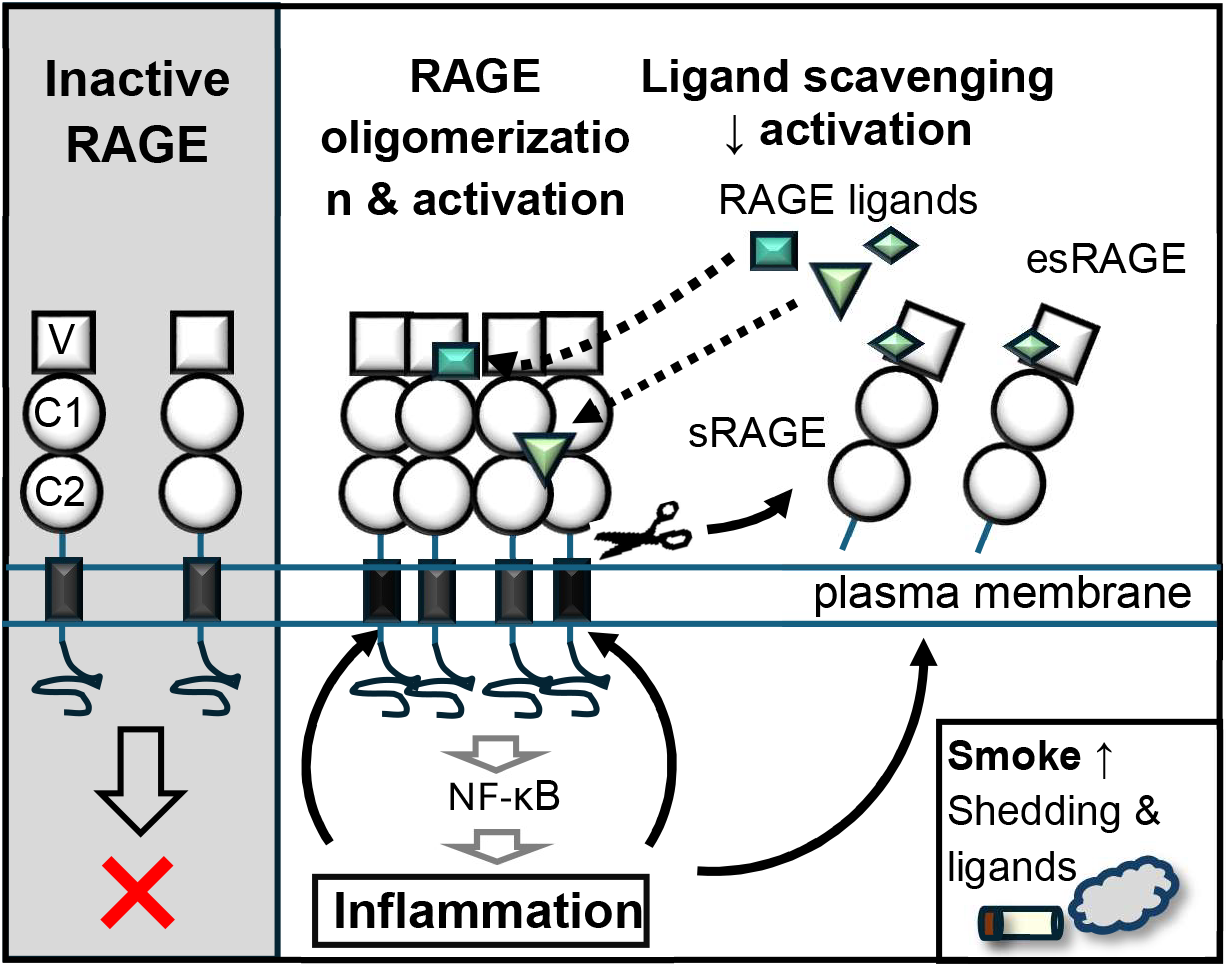
Function of the RAGE protein in cells. RAGE has three extracellular domains (V, C1, and C2), a transmembrane (TM) domains, and a cytoplasmic domain. Ligand binding and oligomerization activate RAGE, which triggers multiple inflammation pathways. Soluble RAGE, which is domains V-C1-C2, has a protective as a ligand scavenger. sRAGE is formed by shedding of the three domains and esRAGE is formed by alternative splicing of *AGER*.

Evaluating the importance of observed gene variants is a challenge and next-generation pathogenicity predictors can be tempting to help. Mutations range from large-scale changes, such as copy number variants, frameshift variants, splicing variants, and gain of STOP variants. Other mutations result in missense variants, which are non-synonymous single nucleotide polymorphisms (SNPs) or variants that change the amino acid type in the encoded protein.

Public databases aggregate human variants, such as ClinVar^12^, which databases human variants involved in disease and drug responses, and the genome aggregation database (gnomAD^13^). gnomAD aggregates large exome and genome sequencing sets as well as providing an estimate in the variant frequencies. There are multiple computational methods of predicting the effects of missense variants. Sequence conservation is a common source of information on protein function for prediction software, since residues associated with protein functions are generally conserved by evolution^14-19^. Incorporation of known protein structures and biophysics adds structural context to possible changes due to missense variants^16-19^. The training algorithms and sets vary the most between softwares.

Mutations can affect the protein expression levels, the stability, the relative amounts of different splicing variants, and the functional efficiency. Clinical databases, such as ClinVar and gnomAD, have observed RAGE variants. New innovations in protein structure prediction and deep-machine learning have enhanced our ability to predict the potential effects of mutations, though the accuracy of these predictions still need to be tested by comparing them to known patient and structural information. As a case study, we present an overview of RAGE variants observed in the ClinVar and gnomAD and evaluate the performance of the AlphaMissense (AM) predictions. The nuances in evaluating predicted pathogenicities are discussed in context with known RAGE structure and mechanism. Additionally, we evaluate the efficiency of a number of prediction algorithms (SIFT, FATHMM, Poly-Phen2, MutPred, AM) in predicting missense mutations associated with COPD-related proteins.

## Materials and Methods

### Case study: RAGE variants

The gnomAD v4.1.0 Genome Aggregation database (https://gnomad.broadinstitute.org/) aggregates large-scale human exome and sequencing data sets^13^ and contains allele frequencies of observed variants. We downloaded the gnomAD *AGER*/RAGE missense, synonymous, frameshift, and truncations (new STOP codon) variants on 06/04/25. A P291L missense cariant was listed. Since wildtype RAGE has I291 and P293, it was unclear if is due to a difference reference sequence. We excluded P291L from the discussion. ClinVar (https://www.ncbi.nlm.nih.gov/clinvar/) includes clinically-observed variants^12^ that were downloads for *AGER*/RAGE on 06/04/25. The AlphaMissense scores for RAGE were downloaded (https://alphamissense.hegelab.org/) as before. The ClinVar data, gnomAD population frequency, and AlphaMissense scores and predictions^19,20^ are listed in **Table 1**.

**Table 1:** *AGER*/RAGE clinical variant information from ClinVar and gnomAD databases and predicted AlphaMissense pathogenicity score (sorted by Allele frequency)

| Protein<br>Consequence | Transcript<br>Consequence | rsID | ClinVar<br>accession<br>number | ClinVar<br>Germline<br>Category<sup>1</sup> | gnomAD<br>Allele<br>frequency<sup>2</sup> | AlphaMissense score<sup>3</sup> |
| --- | --- | --- | --- | --- | --- | --- |
| p.Gly82Ser | c.244G>A | rs2070600 | VCV002582841 | Uncertain | 0.056141623 | 0.3066 |
| p.Arg77Cys | c.229C>T | rs80096349 | VCV000772876 | Benign | 0.00583062 | 0.2295 |
| p.Arg369Gln | c.1106G>A | rs3176931 | VCV000769678 | Benign | 0.003461332 | 0.0703 |
| p.Gly309Arg | c.925G>A | rs138186526 | VCV000782997 | Likely benign | 0.001463366 | 0.0829 |
| p.Trp271Arg | c.811T>A | rs140930365 | VCV000710267 | Conflicting<br>classifications | 0.00105646 | 0.9683 |
| p.Cys301Ser | c.902G>C | rs138178120 | VCV000710266 | Conflicting<br>classifications | 0.001056014 | 0.9587 |
| p.Arg314His | c.941G>A | rs77170610 | VCV000785164 | Benign | 0.001048483 | 0.0714 |
| p.Glu393Lys | c.1177G>A | rs376556772 | VCV002567861 | Uncertain | 0.000100366 | 0.0888 |
| p.Arg198Trp | c.592C>T |  | VCV003275748 | Uncertain | 9.2405E-05 | 0.0694 |
| p.His308= | c.924C>T |  | VCV003842011 | Likely benign | 8.37073E-05 |  |
| p.Cys38Trp | c.114T>G | rs201829223 | VCV002376869 | Uncertain | 8.30835E-05 | 0.9198 |
| p.Val261Asp | c.782T>A | rs201178949 | VCV002392022 | Uncertain | 7.50257E-05 | 0.1187 |
| p.Arg218Gln | c.653G>A | rs766191033 | VCV002314946 | Uncertain | 4.7163E-05 | 0.0911 |
| p.Arg57Trp | c.169C>T | rs766743440 | VCV002351356 | Uncertain | 3.84432E-05 | 0.2359 |
| p.Gly246Asp | c.737G>A | rs150282885 | VCV002388710 | Uncertain | 3.71995E-05 | 0.089 |
| p.Ala355Thr | c.1063G>A |  | VCV003506487 | Uncertain | 3.65682E-05 | 0.0988 |
| p.Arg29Gln | c.86G>A |  | VCV003842020 | Uncertain | 2.85197E-05 | 0.0876 |
| p.Val255Ile | c.763G>A | rs772300820 | VCV002523346 | Uncertain | 2.04594E-05 | 0.0911 |
| p.Arg198Gln | c.593G>A |  | VCV003275758 | Likely benign | 1.79894E-05 | 0.0671 |
| p.Pro202Ser | c.604C>T |  | VCV003275768 | Uncertain | 1.73684E-05 | 0.0641 |
| p.Gly4Arg | c.10G>A | rs769151684 | VCV002603664 | Uncertain | 1.61235E-05 | 0.2051 |
| p.Gly252Ser | c.754G>A | rs766692052 | VCV002225941 | Uncertain | 1.30201E-05 | 0.1234 |
| p.Arg29Trp | c.85C>T |  | VCV003506535 | Uncertain | 1.3019E-05 | 0.1355 |
| p.Pro289Arg | c.866C>G | rs948481571 | VCV003095357 | Uncertain | 1.17857E-05 | 0.0798 |
| p.Gly70Ser | c.208G>A | rs769914792 | VCV002471491 | Uncertain | 1.17793E-05 | 0.0738 |
| p.Gln100Leu | c.299A>T | rs771797821 | VCV003095327 | Uncertain | 9.91984E-06 | 0.1268 |
| p.Arg178Ser | c.534G>C |  | VCV003842039 | Uncertain | 9.2998E-06 | 0.1811 |
| p.Asn389His | c.1165A>C | rs370335494 | VCV002605044 | Uncertain | 9.2926E-06 | 0.0868 |
| p.Ala375Val | c.1124C>T | rs1332735215 | VCV002282320 | Likely benign | 8.05369E-06 | 0.0711 |
| p.Val276Met | c.826G>A | rs764105620 | VCV002377323 | Likely benign | 4.95939E-06 | 0.0693 |
| p.Pro224His | c.671C>A |  | VCV003506498 | Uncertain | 4.34314E-06 | 0.1272 |
| p.Ser74Gly | c.220A>G |  | VCV003275799 | Uncertain | 3.71985E-06 | 0.0894 |
| p.Ser307Ile | c.920G>T | rs1786360349 | VCV002494005 | Uncertain | 3.10031E-06 | 0.1305 |
| p.Arg385Cys | c.1153C>T |  | VCV003506508 | Uncertain | 3.09757E-06 | 0.2306 |
| p.Asn54Ser | c.161A>G | rs890847537 | VCV003095313 | Uncertain | 2.48044E-06 | 0.2139 |
| p.Asn81Ser | c.242A>G | rs1044599245 | VCV002274933 | Uncertain | 1.85986E-06 | 0.095 |
| p.Val63Gly | c.188T>G | rs748270365 | VCV002301361 | Uncertain | 1.85981E-06 | 0.1994 |
| p.Pro293Leu | c.878C>T | rs201967398 | VCV002330677 | Uncertain | 1.85945E-06 | 0.0707 |
| p.Ile291Thr | c.872T>C | rs1194395987 | VCV003095358 | Uncertain | 1.24062E-06 | 0.0625 |
| p.His305Arg | c.914A>G | rs1391153880 | VCV002605889 | Uncertain | 1.2403E-06 | 0.0899 |
| p.Gly56Ser | c.166G>A |  | VCV003275779 | Uncertain | 1.24007E-06 | 0.1861 |
| p.Ile30Val | c.88A>G | rs1034066092 | VCV002359551 | Uncertain | 1.24005E-06 | 0.1338 |
| p.Ala41Val | c.122C>T |  | VCV003842068 | Uncertain | 1.23986E-06 | 0.0904 |
| p.Pro71Thr | c.211C>A |  | VCV003506527 | Uncertain | 1.23984E-06 | 0.0795 |
| p.Pro281Leu | c.842C>T | rs566837079 | VCV002549143 | Uncertain | 6.20227E-07 | 0.075 |
| p.Gly148Arg | c.442G>A |  | VCV003842059 | Uncertain | 6.19931E-07 | 0.5124 |
| p.Val75Met | c.223G>A |  | VCV003506517 | Uncertain | 6.19921E-07 | 0.1483 |
| p.Ile126Ser | c.377T>G |  | VCV003275790 | Uncertain |  | 0.6847 |
| p.Leu159Val | c.475T>G | rs1340132310 | VCV003095329 | Uncertain |  | 0.0856 |
| p.His180Tyr | c.538C>T | rs1369648421 | VCV003095333 | Uncertain |  | 0.1061 |
| p.Glu371Asp | c.1113G>T |  | VCV003506468 | Uncertain |  | 0.1515 |
| p.Glu392Asp | c.1176G>T |  | VCV003842048 | Uncertain |  | 0.1087 |

<sup>1</sup> The full ClinVar Germline Categories (for RAGE) are Benign, Likely benign, Conflicting classifications of pathogenicity, and Uncertain significance. The labels are condensed for space.
<sup>2</sup> Table is sorted by Allele frequency from largest to smallest
<sup>3</sup> The AlphaMissense score thresholds are 0.34 between Likely Benign and Ambiguous and 0.564 between Ambiguous and Likely Pathogenic.

#### Visualization of PTEN models and structures

All protein structures were visualized using PyMOL™ 2.5.2^21^

### Test set and metrics for evaluating prediction quantity

The efficiency of the prediction softwares (SIFT^15^, FATHMM^16^, Poly-Phen2^17^, MutPred2^18^, AlphaMissense^19,20^) were evaluated relative to protein variants observed in ClinVar. ClinVar is a public database of experimentally and clinically observed variants^12^. ClinVar entries for all proteins in **Table 2** were downloaded June and July 2025 (https://www.ncbi.nlm.nih.gov/clinvar/). All entries labeled with the word “benign” (*e.g*., “benign”, “likely benign”, “benign/likely benign”, etc.) were considered benign. All entries including labeled “pathogenic” (*e.g*., “pathogenic”, “likely pathogenic”, “pathogenic/likely pathogenic”) were considered pathogenic. ClinVar entries with other labels (*e.g*., “Uncertain significance”, “Conflicting classification”) were excluded from the analysis.

**Table 2:** Protein analyzed by pathogenicity predictors.

| Proteins analyzed |  |  |
| --- | --- | --- |
| Gene name | Protein name | Uniprot ID |
| AGER | RAGE | Q15109 |
| IL6 | IL-6 | P05231 |
| CXCL8 | IL-8 | P10145 |
| CRP | C-reactive protein | P02741 |
| SFTPD | Surfactant protein D | P35247 |
| APOA1 | Apolipoprotein A1 | P02647 |
| CCL11 | eotaxin-2 | P51671 |
| TNF | TNFα | P01375 |
| SERPINA1 | AAT, alpha-1-antitrypsin | P01009 |
| SFTPA1 | SP-A | Q8IWL2 |
| HSPB1 | HSP27 | P04792 |
| MMP12 | MMP12 | P39900 |
| MMP9 | MMP9 | P14780 |
| BPIFB1 | BPIFB1 | Q8TDL5 |
| IL33 | IL33 | O95760 |
| HMGB1 | HMGB1 | P09429 |
| TLR4 | Toll-like receptor 4 | O00206 |
| LCN1 | Lipocalin 1 | P31025 |
| FGA | Fibronogen alpha chain | P02671 |

Each software has different thresholds and categorization terms. The SIFT webserver for single submitted protein sequences^15^ predicted all 20 possible missense variants at each position (https://sift.bii.a-star.edu.sg/www/SIFT_seq_submit2.html, downloaded July 2025). SIFT scores > 0.05 were considered benign for this analysis and pathogenic otherwise. The UniProt-SwissProt 2010_09 database was searched with a 3.00 median conservation of sequences and sequences with more than 90% identity to the query sequences were excluded. The MutPred2 webserver was utilized to predict variants observed in ClinVar^18^

(http://mutpred2.mutdb.org/index.html, downloaded July 2025). If the MutPred2 score ≤ 0.5 it was considered benign and was considered pathogenic otherwise. The FATHMM (Functional Analysis through Hidden Markov Models, v. 2.3) webserver^16^ focusing of coding variants involved in inherited diseases predicted pathogenicity for observed ClinVar variants (https://fathmm.biocompute.org.uk/inherited.html, downloaded July 2025). FATHMM entries labeled “TOLERATED” were considered benign and those labeled “DAMAGING” were considered pathogenic. The Poly-Phen2 (Polymorphism Phenotyping v. 2) webserver^17^ in batch query mode was utilized to predict the ClinVar observed variants (http://genetics.bwh.harvard.edu/pph2/bgi.shtml, downloaded July 2025). Poly-Phen2 entries labeled “possibly damaging” or “probably damaging” were considered pathogenic and those labeled “benign” were considered benign. AlphaMissense (AM) makes predictions for all 19 possible non-synonymous missense mutations at each protein position. The AlphaMissense scores were downloaded^19,20^ (https://alphamissense.hegelab.org/), July 2025 for each protein in **Table 1**). AlphaMissense has three categories of predictions: “likely benign”, “ambiguous”, and “likely pathogenic”. “Likely benign” were considered benign, “likely pathogenic” was pathogenic, and “ambiguous” entries were excluded from the performance analysis.

The efficiency of the prediction softwares were evaluated for each of the proteins in **Table 2** as binary classifications defining benign as negative and pathogenic as positive. The comparison between the ClinVar labeled variants and the predictions were utilized to calculate the true negative (TN), true positive (TP), false negative (FN), and false positive (FP) rates for each software. Further performance metrics can be done with the following equations (**Table 3**):

**Table 3:** ClinVar data and predictor analytics.

|  | SIFT <sup>4</sup> | FATHMM <sup>5</sup> | PolyPhen-2 <sup>6</sup> | MutPred2 <sup>7</sup> | AlphaMissense <sup>8</sup> |
| --- | --- | --- | --- | --- | --- |
| ClinVar Counts total <sup>1</sup> | 1361 | 1361 | 1361 | 1361 | 1361 |
| ClinVar Counts <sup>2,3</sup> | 216 | 216 | 216 | 216 | 216 |
| Counts benign <sup>2</sup> | 133 | 133 | 133 | 133 | 133 |
| Counts pathogenic <sup>3</sup> | 76 | 76 | 76 | 76 | 76 |
| Counts benign predicted | 109 | 101 | 108 | 161 | 134 |
| Counts pathogenic predicted | 107 | 115 | 108 | 55 | 61 |
| Counts ambiguous/other predicted | -- | -- | -- | -- | 21 |
| Total analyzed | 216 | 216 | 216 | 216 | 195 |
| % analyzed | 15.9% | 15.9% | 15.9% | 15.9% | 14.3% |
| TN (true negative) <sup>9</sup> | 94 | 76 | 99 | 124 | 116 |
| TP (true positive) <sup>9</sup> | 63 | 51 | 68 | 43 | 49 |
| FN (false negative) <sup>9</sup> | 15 | 25 | 9 | 37 | 18 |
| FP (false positive) <sup>9</sup> | 44 | 64 | 40 | 12 | 12 |
| Accuracy | 0.73 | 0.59 | 0.77 | 0.77 | 0.85 |
| Precision (Positive prediction value) | 0.59 | 0.44 | 0.63 | 0.78 | 0.80 |
| Negative prediction value | 0.86 | 0.75 | 0.92 | 0.77 | 0.87 |
| Specificity (True negative rate) | 0.68 | 0.54 | 0.71 | 0.91 | 0.91 |
| Sensitivity (True positive rate) | 0.81 | 0.67 | 0.88 | 0.54 | 0.73 |
| False positive rate | 0.32 | 0.46 | 0.29 | 0.09 | 0.09 |
| MCC | 0.47 | 0.20 | 0.57 | 0.50 | 0.65 |

<sup>1</sup>: ClinVar entries list as "Uncertain significance", "conflicting classification of pathogenicity", and unassigned entries are not used in calculations.
<sup>2</sup>: ClinVar entries listed using the term "benign" (e.g., "benign", "likely benign", "benign/likely benign", etc) were considered Benign for this analysis.
<sup>3</sup>: ClinVar entries listed using the term pathogenic (e.g., "pathogenic", "likely pathogenic", "pathogenic/likely pathogenic", etc) were considered pathogenic for this analysis.
<sup>4</sup>: SIFT scores > 0.05 were considered benign; otherwise, as pathogenic.
<sup>5</sup>: FATHMM entries labeled "TOLERATED" were considered benign and "DAMAGING" as pathogenic.
<sup>6</sup>: Poly-Phen2 entries labeled as "possibly damaging" or "probably damaging" were considered pathogenic. "benign" entries were benign.
<sup>7</sup>: If MutPred2 score ≤ 0.5, then entry classified as benign; otherwise as pathogenic.
<sup>8</sup>: Ambiguous AlphaMissense predictions were not used in calculations. AlphaMissense also did not analysis Met1 residues.
<sup>9</sup>: Benign variants are considered negative and pathogenic are positive.

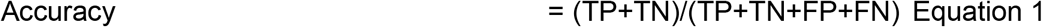

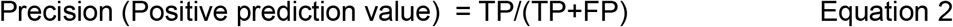

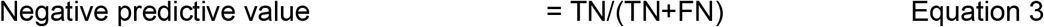

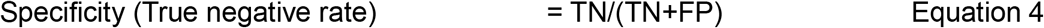

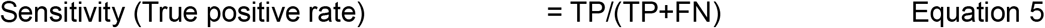

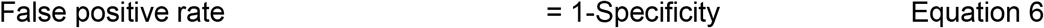

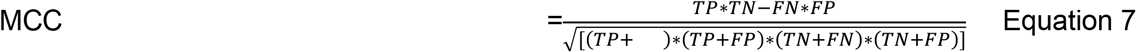

## Results

### Case study: RAGE protein and utilizing structural biology to understand variant predictions

There are national consensus guidelines for how to categorize the phenotypic consequences from variants and how strong different types of evidence are in this categorization. The American College of Medical Genetics and Genomics (ACMG) and the Association of Molecular Pathology (AMP) published a guideline utilizing five categories: pathogenic (i.e. disease-causing), likely pathogenic, uncertain significance, likely benign, and benign^25^. There are other national guidelines, such as from the UK Association of Clinical Genomic Sciences (ACGS), and refinements of the 2015 guidelines by the US ClinGen Sequence Variant Interpretation (SVI) working group^26,27^. ClinVar utilizes the five categories from the ACMG/AMP 2015 guidelines, as well as a Conflicting Reports category.

Missense variants result in a change of amino acid type in the RAGE protein. RAGE missense variants listed in the public database ClinVar^12^ are mostly classified as being of “Uncertain significance” (**Table 1**). Two variants, W271R and C301S have “Conflicting reports of pathogenicity”. The remaining variants have likely benign or benign consequences. The Genomic Aggregation database (gnomAD, v.4.1.0) aggregates and summarizes large-scale human exome and sequencing data sets and is meant to give an indication of the baseline variability in the human genome, including estimates of the allele frequency of a particular variant. The most common missense mutation is G82S, with a frequency of 0.056 (**Table 1, Figure 2A**). This is an order-of-magnitude more common that the next most common mutations—R77C, W271R, C301S, G309R, R314H, and R369Q. These variants are more common than the rest of the missense, truncation (gain of STOP codon), and frameshift mutations.

**Figure 2.**
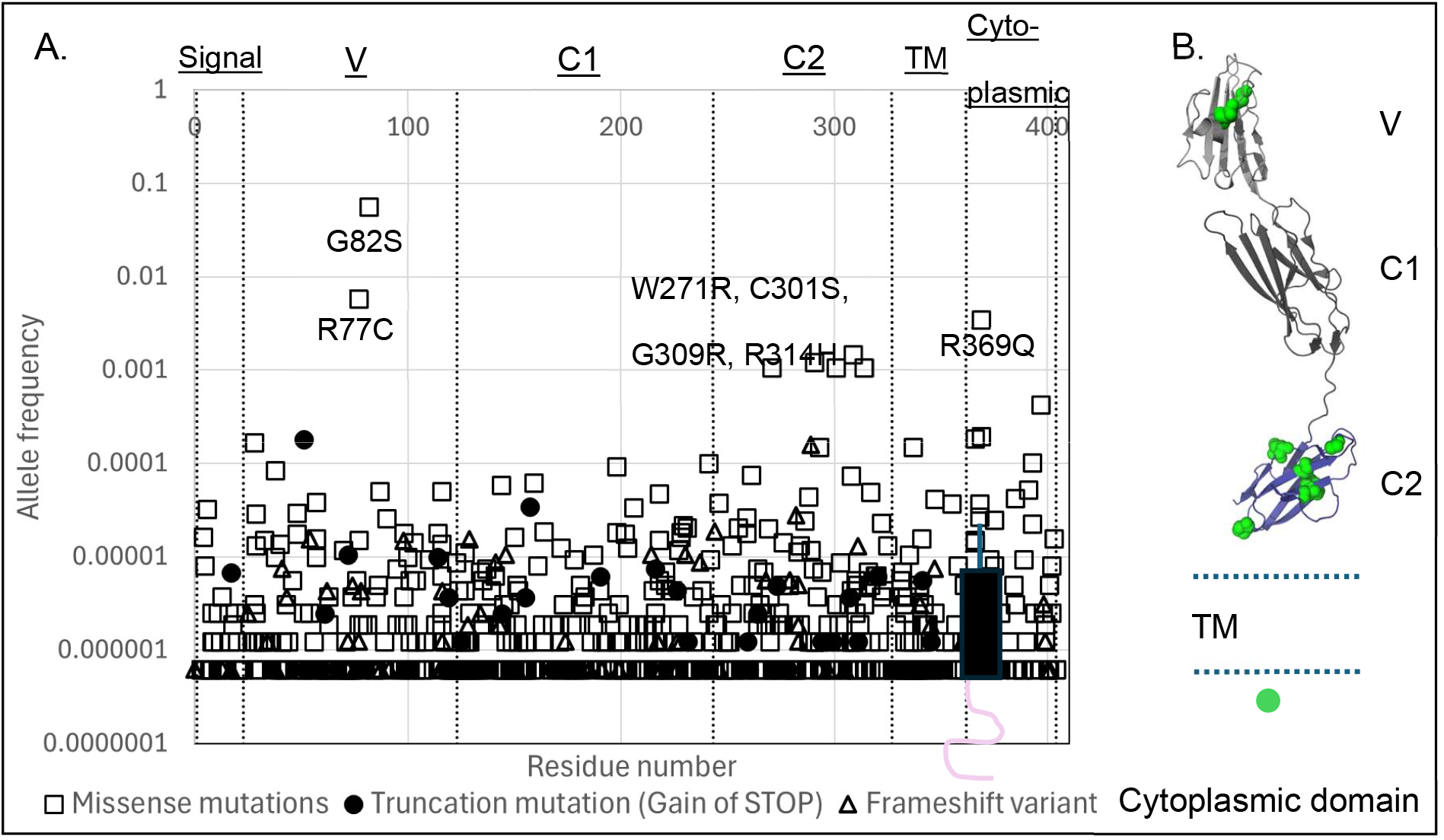
A. Frequency of RAGE missense variants. (marked as *open squares*), truncation (as *circles*), and frameshift variants (as *open triangles*). The gnomAD allele frequency plotted versus the residue number with the boundaries between domains indicated as *dashed lines*. B. Location of the most common missense mutations. The most common mutations, based on gnomAD allele frequency, are mapped *in green* onto the cartoon of RAGE. Structures of domains V, C1, and C2 are from X-ray structure of soluble RAGE (PDBID 4LP5).

The position of missense variants can potentially have consequences to the protein. The V, C1, and C2 domains are outside the cell in the full-length RAGE receptor and are included in the soluble, truncated sRAGE (**Figure 1**). G82S and R77C variants are in the V domain (**Figure 2A and 2B**), which is the major ligand binding site for the protein^28-32^. ClinVar classifies R77C as benign and G82S as “uncertain significance” (**Table 1**). Generally, variants with observed frequencies of >5% are considered benign^25^. However, G82S is both a common human polymorphism of RAGE (>5% in gnomAD) and has important physiological consequences. W271R, C301S, G309R, and R314H are present in the C2 domain. The V, C1, and C2 domains each have a disulfide bond^4,5,28,41,42^. C301 is part of the C2 domain disulfide bond, which is disrupted in the C301S variant. W271R mutates a core residue of the domain. These two have “conflicting classifications of pathogenicity” in ClinVar. ClinVar classifies G309R as likely benign and R314H as benign. R369Q is in the cytoplasmic domain, though its position near the boundary with the end of the transmembrane domain, as well as its positive charge, suggests that it interacts with the negatively-charge leaflet of the plasma membrane^43^.

### Case study: Predicting RAGE missense variants

Computational pathogenicity predictions are usually considered supportive to a variant’s categorization if multiple independent predictions are made^25^, though some recent efforts has been made to discover which software can be considered supporter evidence for categorization^27^. The AlphaMissense (AM) deep-machine learning algorithm is trained on sequence and structural data, as well as population frequency data^19,20^. It predicts all possible amino acid substitutions at each position in the protein (19 substitutions* 404 residues = 7,676 predicted missense mutations for RAGE). It predicts if any mutant is “Likely benign”, Ambiguous”, or “Likely pathogenic”. Some positions in a protein are known to be more accommodating to mutations, particularly surface, loop, and disordered regions, while parts of the dense core of the protein, binding sites, and the catalytic residues are generally more sensitive. To get a feel for how sensitive a position is to any kind of mutation, we summed the number of missense mutations that were predicted “Likely pathogenic” at each position (**Figure 3A**). As expected, the surface residues tend to be more accommodating of mutations while core residues are predicted to be more sensitive (**Figures 3A and 3B**). Structural features, like Cys-Cys disulfide bonds, are also known to be more sensitive to mutations.

**Figure 3.**
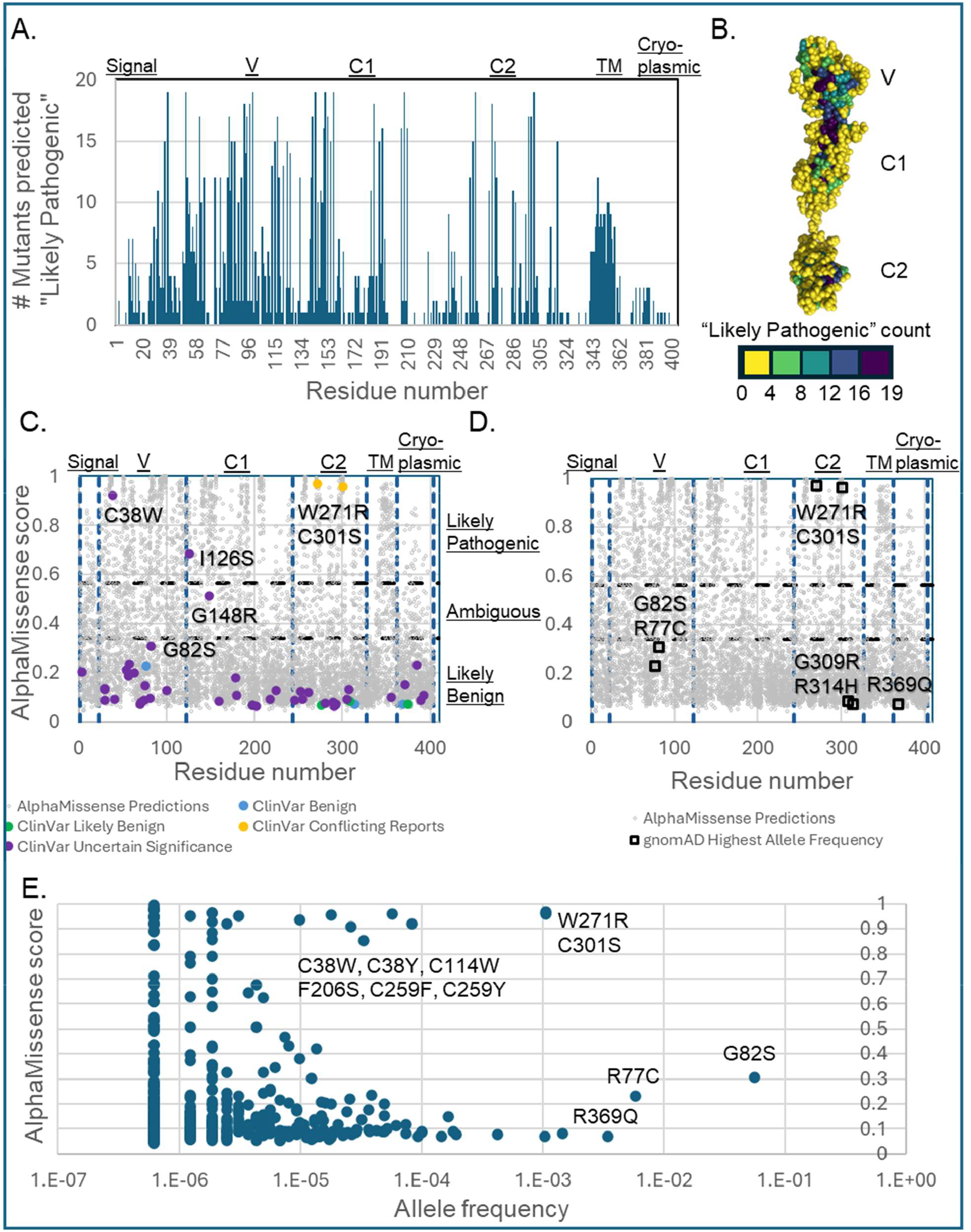
AlphaMissense (AM) predictions of RAGE mutation pathogenicity. A. Sensitivity of RAGE residues to mutations. Number of “Likely Pathogenic” mutations predicted by AM plotted versus residue numbers. AM predicted the 19 possible animo acid substitutions for each residue position, so a position with a higher number of pathogenic predictions may be more sensitive to mutation. Portions of the V, C1, C2, and TM domains are more sensitive than the more flexible signal peptide and cytoplasmic domain. **B**. Mapping sensitivity onto sRAGE structure (PDBID 4LP5), shown in sphere representation). Surface, linker, and loop residues have few predicted pathogenic mutations (count=0 to 3, shown *in yellow*). Internal core residues tend to be most sensitive (*in purple*). Positions where mutations have more intermediate effects (count=4 to 15) are *in green to blue*. **C**. AlphaMissense predicts most of the ClinVar RAGE mutations as benign. AM scores for each possible mutation is shown in grey and clinically-observed RAGE mutations are indicated for ClinVar entries labeled “Benign” (*in blue*), “Likely benign” (*in green*), “Conflicting classifications of pathogenicity” (*in orange*), and “Uncertain significance” (*in purple*). The two “Conflicting” entries, W271R and C301S, are predicted “Likely pathogenic”. The “Uncertain” mutations C38W, I126S, and G148R are predicted to be “likely pathogenic’ or ambiguous. Notably, G82S, which has known clinical consequences, is predicted to be “likely benign”. **D**. Most common missense mutations are predicted to be “likely benign”, except for W271R and C301S. The missense mutations with the greatest gnonAD allele frequency are indicated as *open squares*. E. Plotting allele frequency versus AM score reveals other potentially pathogenic mutations. C38W, C38Y, C114W, F206S, C259F, and C259Y are predicted “Likely pathogenic” and have 10^-4^ to 10^-5^ allele frequency.

AM predicts most of the ClinVar RAGE mutations as benign (**Figure 3C, Table 3**). AM scores for each possible mutation is shown in grey and clinically-observed RAGE mutations are indicated for ClinVar entries labeled “Benign” (*in blue*), “Likely benign” (*in green*), “Conflicting classifications of pathogenicity” (*in orange*), and “Uncertain significance” (*in purple*). The two “Conflicting” entries, W271R and C301S, are predicted “Likely pathogenic”. The “Uncertain” mutations C38W, I126S, and G148R are predicted to be “likely pathogenic’ or ambiguous. Notably, G82S, which has known clinical consequences, is predicted to be “likely benign”. The G82S polymorphism, which is located in the V domain, increases ligand affinity^44,45^, while decreasing protein stability^46^. Most common missense mutations are predicted to be “likely benign”, except for W271R and C301S (**Figure 3D**). The missense mutations with the greatest gnonAD allele frequency are indicated as *open squares*. Notably, plotting allele frequency versus AM score reveals other potentially pathogenic mutations (**Figure 3E**). C38W, C38Y, C114W, F206S, C259F, and C259Y are predicted “Likely pathogenic” and have 10^-4^ to 10^-5^ allele frequency.

### Evaluation of pathogenicity predictors

Variant pathogenicity can be predicted utilizing tools, such as SIFT, FATHMM, Poly-Pred2, MutPred2, and AM^14-19,47^. Similar analyses of prediction softwares sampling variants from the entire human proteome or subsets of proteins related to specific diseases^20,27,48,49^. SIFT, FATHMM, MutPred2,and Poly-Phen2 are considered by specifically in classification protocol literature^25,27^, but AM is newer. Here, we utilized a selected set of COPD-related proteins (**Table 2**) to test the quality of their predictions. As a comparison, we used ClinVar, a public database of clinically and experimentally observed variants, extracting the missense variants. We excluded the synonymous variants whose nucleotide changes do not result in changes in protein amino acids. Variant pathogenicity prediction relies on different algorithms. Sequence conservation is a common source of information on protein function for the SIFT, FATHMM, Poly-Phen2, MutPred2, and AM prediction software, since residues associated with protein functions are generally conserved by evolution^14-19^. Incorporation of known protein structures and biophysics adds structural context to possible changes due to missense variants^16-19^. The training algorithms and sets vary the most between softwares. FATHMM utilizes hidden Markov models^16^. MutPred2, Poly-Phen2, and AM use probabilistic or machine-learning algorithms^17-19^. SIFT makes predictions based of sequence homology, noting that residues associated with protein functions are conserved by evolution^14,15^. FATHMM (functional analysis through hidden Markov models) also rely on multiple sequence alignments and known conserved domains and utilize hidden Markov models^16^. AM is trained on sequence, predicted structures form AlphaFold^50,51^, and weak labels from population frequency data and makes predictions on the pathogenicity of a specific missense variant^19,20^.

We evaluated the performance of these predictions as binary classifications. Benign variants were considered negative and pathogenic variants as positive. ClinVar has multiple categories. All entries labeled “benign”, “likely benign”, or “benign/likely benign” were considered negative. All entries labeled “pathogenic”, “likely pathogenic”, or “pathogenic/likely pathogenic” were considered positive. ClinVar entries with other labels (e.g., “Uncertain significance”, “Conflicting classification”) were excluded from the analysis. Of the 1361 ClinVar non-synonymous missense variants for the 19 COPD-related proteins (**Table 2**), 216 were as either benign (negative) or pathogenic (positive) (**Table 3**). Each prediction method has their own threshold for determining if a variant is benign or pathogenic (see Materials and Methods). AM also has a third category “ambiguous” that was excluded from analysis. Additionally, AM does not predict variant for the first Met1 residue of each protein sequence. Therefore, the 216 variants were analyzed for all algorithms except AM, which has 195 (**Table 3**). The observational ClinVar data and predictions were utilized to calculate the true negative (TN), true positive (TP), false negative (FN), and false positive (FP) counts for each prediction method. We evaluated the performance of these predictions (see Materials and Methods). The accuracy is the fraction of true predictions, both positive and negative. All methods except FATHMM had greater than 0.7 accuracy with AM being the most accurate with 0.85 (**Table 3**). Precision is the positive predictive value and is the fraction of positive predictions that were true. The AM predictions had the greatest precision. Negative predictive value is greatest for Poly-Phen2, followed by AM and SIFT. The specificity is the true negative rate and the sensitivity is the true positive rate. AM and MutPred2 have the highest specificity and Poly-Phen2, then SIFT and AM, have the greatest sensitivity. The false positive rate is the lowest for AM and MutPred2. The Matthews correlation coefficient (MCC) is a quality metric for the binary classifications^52^. MCC evaluates the performance for true negative, true positive, false negative, and false positive predictions and does not require equal proportions of positive and negative entries. The MCC for AM is 0.65 with an accuracy of 0.85 and false positive rate of 0.09. This means that AM predicted about 85% of the variants classified as either benign or pathogenic in ClinVar correctly. It should be noted that the ClinVar labels were assumed to be correct and only the variants with benign or pathogenic classifications in both ClinVar and AM were analyzed. The binary classification statistics may not be representative of the more ambiguously classified variants.

## Discussion

There are public databases of observed variants for the general and clinical populations, as well as new tools for prediction of the effects of these variants, though interpretation of these data can be challenging. The G82S RAGE variant is a common human polymorphism with >5% allele frequency in the Genomic Aggregation (gnomAD) database^13^. Generally, variants with observed frequencies of >5% are considered benign^25^. Here, notably, G82S was predicted to be “likely benign” by AM (**Figure 3C, Table 3)**, the best performing of the five tested predictors (**Table 3**). However, G82S has important physiological consequences. It has known protective effects for COPD^2,33-38^, but deleterious consequences for Alzheimer’s disease^39^ and cardiovascular disease^40^. G82S is in the RAGE V domain, which is a major binding interface for DAMP ligands that trigger inflammation pathways via the RAGE receptor^44,53^ (**Figure 1**). G82S increases binding affinities for these ligands. The variant also increases the binding affinities in sRAGE, which scavenges the ligands, counteracting the inflammation. The mutation site is adjacent to the N81 N-linked glycosylation site. In wildtype RAGE, N-linked glycosylation decreases binding affinity^44^. In G82S RAGE, while the level of glycosylation at N81 is increased^44,45^, the binding affinity is also increased^45^. This suggests that G82S may interfere with the glycosylation-based regulation of ligand binding. There are multiple forms of RAGE in the cells, which can be differentially affected by a polymorphism. G82S increases the amount of full-length RAGE receptor^54^ relative to wildtype RAGE, but notably reduces the amount of soluble sRAGE^1,2^.

Our analysis of public clinical databases (ClinVar, gnomAD), as well as the pathogenicity predictions of AlphaMissense highlighted several variants that would disrupt the disulfide bonds in the RAGE V, C1, and C2 domains. In a disulfide bond, the sidechain sulfur atoms of two cysteine residues form a S-S covalent bond. Extracellular spaces between cells and the ER of eukaryotic cells are non-reducing and do not disrupt disulfide bonds, while reducing environments contain enzymes that break these bonds. These bonds can be structural, particularly for domains exposed to the extracellular space like RAGE V, C1, and C2 domains. Each of these domains have one disulfide bond: C38-C99 in V domain, C144-C208 in C1, and C259-C301 in C2^4,5,28,41,42^. RAGE associates with other copies of itself (oligomerizes), both in its *apo* (unbound) form and while ligand-bound (**Figures 1 & 2**). Activation of the inflammatory response was closely linked to RAGE receptor’s oligomeric state^4,6,55,56^. FRET and chemical crosslinking studies demonstrated that RAGE oligomerizes on the plasma membrane (PM) in HEK293T cells^46,57^. RAGE was observed as a distribution of monomer to tetramer in micelles via native mass spectrometry (MS)^8^. Oligomerization involves the multiple RAGE domains: inter-subunit interactions between the V and C1 domains (observed via crosslinking MS (XL-MS))^8^, suggested disulfide bonding between C2 subunits^7,58^, and between the C1-C2 linkers^59^.

The performance of pathogenicity predictors have made remarkable improvements (**Table 3)**. National consensus guidelines on how to categorize the phenotypic consequences from variants generally consider *in silico* predictions to be supportive of a variant’s classification if there are multiple independent results^25^. More recent refinements of the 2015 guidelines by the US ClinGen Sequence Variant Interpretation (SVI) working group suggest that some prediction methods be given more weight in classification ^26,27^. However, care does need to be taken when interpreting the prediction results, with knowledge of the structural biology and the literature taken into consideration.

## Author Approvals

All authors have seen and approved the manuscript and it has not been accepted or published elsewhere.

## Competing interests

There are no competing interests.

## Funding information

This study was funding in part by the COPDGene study (<u>NCT00608764</u>) and Cleveland Clinic Catalyst grant IF111938.

## Notes

### Competing Interest Statement

The authors have declared no competing interest.

